# Transcriptome Data Analysis of Primary Cardiomyopathies Reveals Perturbations in Arachidonic Acid Metabolism

**DOI:** 10.1101/2022.11.06.515334

**Authors:** Pankaj Kumar Chauhan, R. Sowdhamini

## Abstract

Cardiomyopathies are complex heart diseases with significant prevalence around the world. Among these, primary forms are the major contributor to heart failure and sudden cardiac death. As the high-energy demanding engine, the heart utilizes fatty acids, glucose, amino acid, lactate and ketone bodies as energy to meet its requirement. However, continuous myocardial stress and cardiomyopathies drive towards metabolic impairment that advances heart failure (HF) pathogenesis. So far, metabolic profile correlation across different cardiomyopathies remains poorly understood. In this study, we systematically explore metabolic differences amongst primary cardiomyopathies. By assessing the metabolic gene expression of all primary cardiomyopathies, we highlight the significantly shared and distinct metabolic pathways that may represent specialized adaptations to unique cellular demands. We utilized publicly available RNA-seq datasets to profile global changes in the above diseases and performed Gene set analysis (GSA). Our analysis demonstrates genes in arachidonic acid metabolism (AA) as significantly perturbated across cardiomyopathy. In particular, arachidonic acid metabolism gene *PLA2G2A* interacts with fibroblast marker genes and can potentially influence fibrosis during cardiomyopathy.

## Introduction

Primary cardiomyopathies, predominantly hypertrophic cardiomyopathy (HCM), dilated cardiomyopathy (DCM), restrictive cardiomyopathy (RCM) and arrhythmogenic cardiomyopathy (ACM) are a growing global burden on public health (1–6). Primary cardiomyopathies, on the whole, are genetic but tend to be influenced by environment and lifestyle (6–12). These heterogeneous heart muscle diseases lead to heart failure and sudden cardiac death (1,6). Notably, metabolic impairment is vital in heart failure (13,14). Generally, the heart meets its energy demand by utilizing fatty acids, glucose, amino acid, lactate and ketone bodies, but cardiomyopathies lead to severe metabolic perturbation (15–17). In cardiovascular research, metabolic explorations have provided new insights (18,19). The role of metabolic genes and pathways has been explored in several cardiomyopathy studies to understand the pathophysiological changes involved in the progression of cardiomyopathy (15–17,20–22). These studies have reported a metabolic shift in energy sources during disease progression. HCM-focused transcriptome analysis showed down-regulated fatty acid metabolism (23). Similarly, in a DCM transcriptome analysis, mitochondrial dysfunction and oxidative phosphorylation pathways were significantly perturbed (24). Concurrently, multi-omics technologies have provided an opportunity to explore metabolic perturbation at a large scale. RNA-seq technique allows rapid measurement of global gene expression in the disease of interest. Further, it provides an indirect mechanism to assess metabolic perturbations (25– 27). Several computational algorithms incorporating transcriptome data and metabolic networks have been developed to assess the perturbation of biological pathways (28–30).

Over the years, oxidative phosphorylation, glucose and fatty acid metabolism have been highlighted to be perturbed in individual cardiomyopathy studies. Albeit being important, individual cardiomyopathy-focused metabolic studies miss overall molecular patterns across cardiomyopathies. A comparative interpretation of the primary cardiomyopathies’ metabolic alterations is crucial for a comprehensive understanding of the mechanisms of metabolic perturbations. To the best of our knowledge, our study is the first exploration that systematically explores metabolic correlations amongst primary cardiomyopathies (HCM, DCM, RCM and ACM).

This study aimed to identify shared metabolic perturbations across cardiomyopathies. We utilized gene expression profiles of primary cardiomyopathies: HCM, DCM, RCM and ACM, and donor samples. We carried out differential gene expression analysis of cardiomyopathy datasets. Further, we performed GSA on each dataset to investigate the metabolic alterations comprehensively. Apart from glycolysis and oxidative phosphorylation, the arachidonic acid metabolism (AA) pathway was significantly altered in all cardiomyopathy sets. Subsequently, we inferred the potential cell types in the candidate pathway using the snRNA-seq dataset. This analysis helped in the identification of marker genes for each cell type and the potential link between AA metabolism genes and cell type marker genes.

Earlier studies did not account for cross-comparison nor focused on arachidonic acid metabolism in depth. AA is a free fatty acid metabolized by cyclooxygenase, lipoxygenase, and cytochrome P450 monooxygenase enzymes into biologically active fatty acid mediators (31). Through these mediators, AA participates in complex cardiovascular functions, including fibrosis (32). Our analysis shows that AA enzymes are expressed in the fibroblasts, cardiomyocytes, smooth muscle cells, monocytes, macrophages and mast cells in the heart tissue. It is relevant to look into the role of AA metabolism in cardiomyopathies. Overall, our study highlights novel and clinically valuable aspects of cardiomyopathies, with implications ranging from prognosis to therapeutic intervention.

## Materials and Methods

### Data Acquisition and Preprocessing

Transcriptomic raw data comprising RNA-seq, single-nucleus RNA-seq (snRNA-seq) and microarray for cardiomyopathy studies were acquired from the European Nucleotide Archive (ENA) database (https://www.ebi.ac.uk/ena/browser/) and Gene Expression Omnibus (GEO) database (https://www.ncbi.nlm.nih.gov/geo/). Search terms “hypertrophic cardiomyopathy”, “dilated cardiomyopathy”, “restrictive cardiomyopathy”, and “arrhythmogenic cardiomyopathy” were searched against these databases to obtain the RNA-seq and microarray dataset results. Each result was manually reviewed and considered for inclusion if (1) the disease samples in the study indicated heart tissues from the cardiomyopathy patients; (2) control samples came from non-failing heart patients. We focused on the studies that were supported in our R-analysis pipeline. We considered seven publicly available cardiomyopathy transcriptomic data, including five RNA-seq (SRP125284, SRP125595, SRP052978, SRP186138 and SRP061888) and two microarray datasets (GSE29819, GSE36961) using these inclusion criteria. One single-nucleus RNA sequencing (snRNA-seq) dataset (GSE183852) with dilated cardiomyopathy (DCM) was selected to explore cell type expression of candidate genes due to the cell annotation in the supplementary files of the original dataset (33). The details of the transcriptome datasets are shown in Supplementary Table **S1**.

### Data Processing and Differential Expressed Genes (DEGs)

Previously described and curated raw data were downloaded and reprocessed to ensure uniform processing and normalization of each study. RNA-seq and microarray studies were processed using independent pipelines. In the RNA-seq pipeline, custom shell scripts were used to download data. Salmon (v1.5.2) (a fast and bias-aware quantification tool) was utilized to align and quantify samples using NCBI human reference transcriptome (Gencode v38) (34). The count data output of Salmon quantification was used for differential gene expression analysis between disease and donor samples. This analysis was performed using the DESeq2 (v1.26.0) package in R (v 3.6.3) (35). It uses the negative binomial distribution to model a statistical analysis for differential gene expression. It also normalizes samples automatically. Wald t-test was applied to the distribution (36). To control the false discovery rate (FDR), the resultant *p-values* were adjusted using Benjamini and Hochberg’s test (BH) correction (37). Genes with *adjusted P* < 0.1 and *|log_2_FC|* ≥ 0.28 were assigned as being differentially expressed. Likewise, microarray datasets were processed with the limma (v3.42.2) package (38). A linear model was constructed between disease and control samples, and the empirical Bayes statistical method was utilized to obtain the significant genes. BH correction was used to obtain the *adjusted P* value. The snRNA-seq expression stored as the R object (33) was processed and analyzed using the R package “Seurat” (version 3.2.3).

### Sample Variability and Study Consistency

Principal component analysis (PCA) and t-distributed stochastic neighbor embedding (t-SNE) were applied to assess the sample variability across datasets. Only RNA-seq datasets were explored for testing. These datasets were normalized using the “DESeq” method in the R package. The R function “prcomp” was used to perform PCA. On the other hand, the “Rtsne” function was utilized to perform t-SNE. Principal component &38; t-SNE plots, Venn diagrams, and heatmap plots were prepared using the ggplot2 R package (version 3.3.5) and Matplotlib package in Python 3.

### Gene Set Analysis (GSA)

The gene set analysis was carried out on the gene expression to identify significantly perturbed pathways in each study. We utilized *p-values* from the differential expression analysis for all genes in individual datasets for this analysis. A ranking was generated based on these *p-values*, representing the Gene level statistics (gene expression). The Kyoto Encyclopedia of Genes and Genomes (KEGG) pathways list file was downloaded from KEGG (https://www.kegg.jp/kegg-bin/get_htext#C1). Parametric analysis of gene set enrichment (PAGE) method was employed for GSA analysis using the Piano (version 2.2.0) package in R (39,40). The PAGE uses the mean of the gene-level statistics of a gene set (a particular pathway in this case) and corrects for the background, represented by all gene-level statistics. The cumulative normal distribution is used to estimate the PAGE gene-level statistics significance. Heatmap of pathways, volcano plots and bar charts were plotted using Matplotlib and Seaborn packages in Python 3.

### Identification and Screening of Significant Pathways

Pathways with the PAGE statistics *p-values* < 0.1 were considered significant. Among many perturbed pathways like glycolysis and oxidative phosphorylation, arachidonic acid metabolism was chosen as the candidate for further analysis due to its significance in all primary cardiomyopathies. To show that the arachidonic acid (AA) metabolism perturbation was consistent in all datasets rather than only individual studies, we used microarray to validate its role in ACM, DCM and HCM independently. We could not find any microarray dataset for RCM, so it was not considered. Gene set analysis of these microarray studies was performed using the sorted Gene level statistics GSA as previously described. Genes differentially regulated in the AA metabolism were mapped to the KEGG mapper using the online server: Pathview Web (https://pathview.uncc.edu/) (41).

### Finding DEGs, Cell types Annotation in Arachidonic Acid Metabolism and Cell-Specific Marker Genes

Finally, to explore AA metabolism genes, AA genes were screened with a cut-off value of *adjusted P* < 0.1 and *|log_2_FC|* ≥ 0.28. To understand the expression of these genes in the different cell types of the heart tissue, gene expression and cell phenotypes were considered from an snRNA-seq study (GSE183852) (33). In this, control samples were filtered to identify the expression of screened AA metabolism genes in the donor heart cell types. Further, marker genes were identified in each cell type. The ‘FindMarkers’ function in the Seurat package was used to accomplish this task with the cut-off values of *adjusted P* < 0.1, minimal percentage > 0.1, and *log_2_FC* ≥ 0.58. These marker genes were utilized to select the cell types significantly enriched within cardiomyopathies DEGs. The GSA method PAGE was employed to evaluate the significance of the DEGs against the cell-specific marker genes in the Piano (version 2.2.0) package in R (39,40). Functional enrichment of cell-specific marker DEGs was performed using online tool ‘g:Profiler’ (https://biit.cs.ut.ee/gprofiler/) (42).

### Identification of Dysregulated Marker Genes Interactions in The Cell Types

For human interactome data, PPI data (HPRD, MINT, IntAct) along with protein complex and kinase substrate data (CORUM, Phosphositeplus) were obtained from our previous study (43). Then, the AA metabolism genes and cell type marker genes were mapped to the human interactome, and a subnetwork consisting of these genes was constructed. Network visualization was executed in Gephi (version 0.9.3).

## Results

### Gene Expression Profile in Primary Cardiomyopathies

We identified and selected seven studies fitting our inclusion criteria (see Table S1 and Methods), consisting of arrhythmogenic cardiomyopathy, dilated cardiomyopathy, hypertrophic cardiomyopathy and restrictive cardiomyopathy samples. Analyses were performed using 4 RNA-seq, one single-nucleus RNA-seq (snRNA-seq) and two microarray datasets. We performed sample variability analysis on normalized RNA-seq studies. The PCA and t-SNE analysis revealed variability in the samples of different datasets. Attributes like demography, genetic differences and tissue biopsies influence the above variability (Figure 1A, B). Since each dataset independently consists of control and disease samples, these variations can be ignored. For the RNA-seq differential gene expression, the negative binomial generalized linear model was used (see Methods). We employed the empirical Bayes method on the generalized linear model for the microarray datasets DEGs. This analysis led to the identification of 4158, 5822, 3048 and 1655 DEGs in ACM, DCM, HCM and RCM RNA-seq studies. We screened KEGG metabolic genes amongst these DEGs and found 527, 644, 346 and 150 differentially regulated metabolic genes (see Figure 1C). Gene expression results indicate that DCM gene expression differed from other primary cardiomyopathies (see Figure 1D).

**Figure 1:**
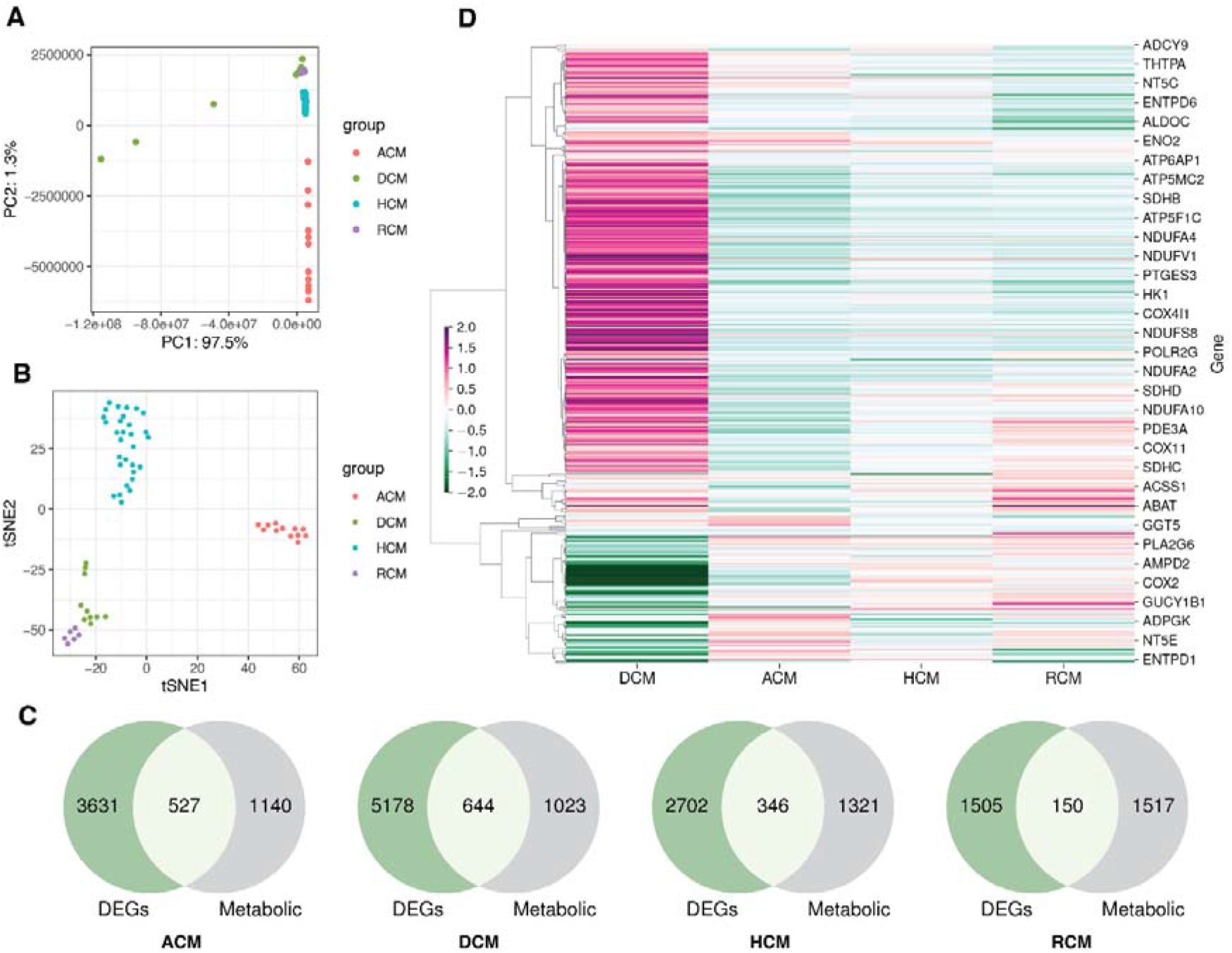
The transcriptome datasets in primary cardiomyopathies. (A) PCA plot of disease and donor samples in ACM, DCM, HCM and RCM. (B) the t-SNE plot disease and donor samples in ACM, DCM, HCM and RCM. (C) Venn diagrams showing DEGs and metabolic genes in each cardiomyopathy. (D) A heatmap showing the metabolic genes expression in ACM, DCM, HCM and RCM phenotypes.

### The Metabolic Pathways are Significantly Dis-regulated in Cardiomyopathies

To further understand the role of metabolic genes in cardiomyopathies, we performed gene set analysis (GSA) on RNA-seq and microarray studies. The *p-values* from the differential expression analysis of each dataset were ranked to estimate the gene level statistics (see Methods). The KEGG pathway metabolic gene signature was used to identify the dis-regulated pathways. The GSA analysis revealed widespread alterations of glycolysis/gluconeogenesis, TCA cycle, oxidative phosphorylation, riboflavin, thiamine, purine and arachidonic acid metabolism (see Figure 2).

**Figure 2:**
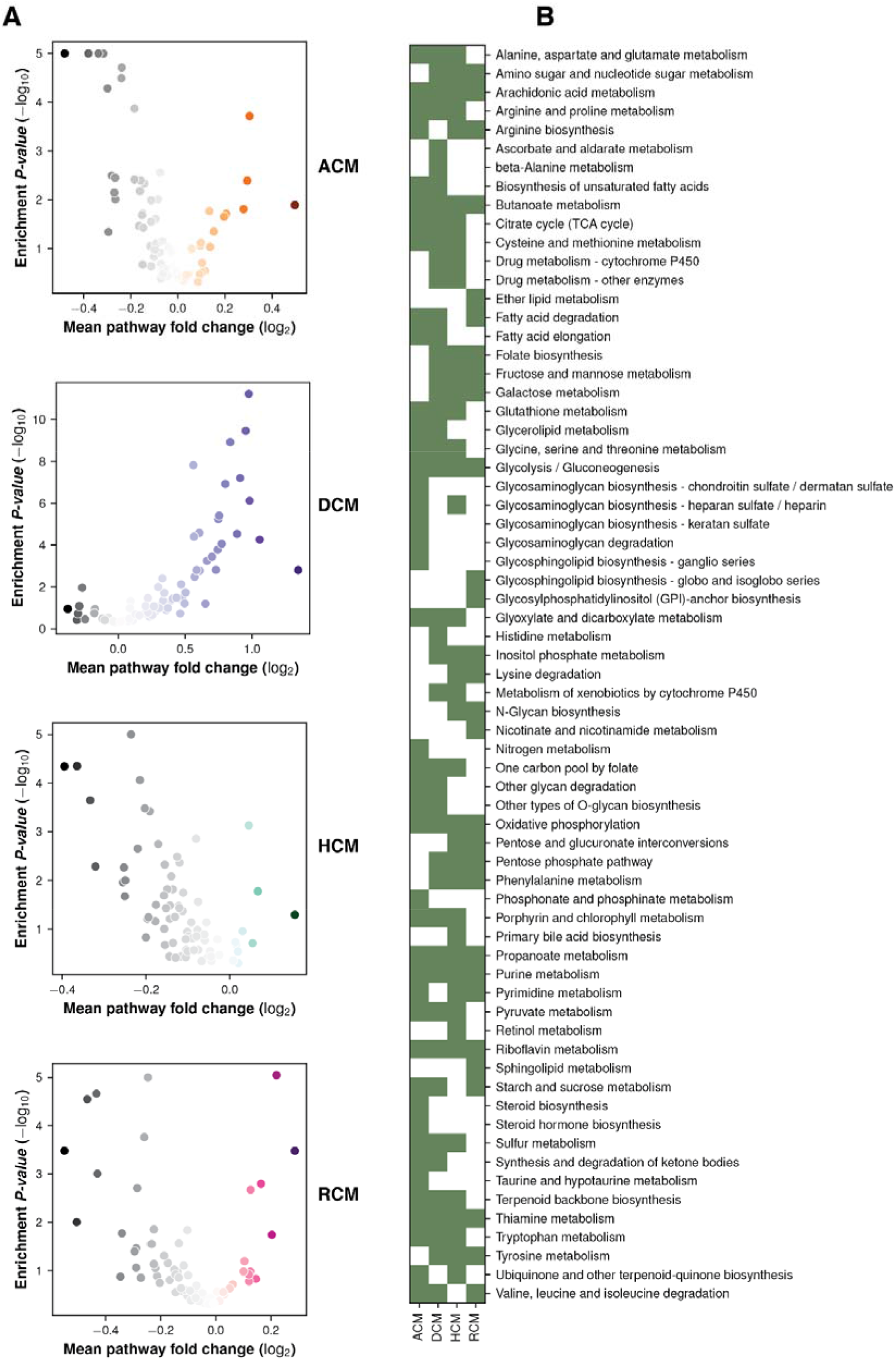
GSA enrichment of KEGG pathways in primary cardiomyopathies. (A) Volcano plots showing pathways enrichment and fold change in primary cardiomyopathies. (B) A heatmap showing major pathways perturbation in ACM, DCM, HCM and RCM phenotypes.

Oxidative phosphorylation metabolism, a mitochondrial-associated process, was significantly downregulated in the ACM, HCM and RCM but not in the DCM. It was upregulated in the DCM phenotype. Similarly, another primary energy source, glycolysis/gluconeogenesis metabolism, was also downregulated in ACM, HCM and RCM and upregulated in DCM. Nucleotide-specific purine metabolism was downregulated in all phenotypes except DCM. In addition, the pathway related to the amino acid arginine biosynthesis was significantly upregulated in the DCM but significantly downregulated in other primary cardiomyopathies. Process related to co-factor metabolism, riboflavin was upregulated in DCM but downregulated in ACM, HCM and RCM. Interestingly, inositol phosphate metabolism was upregulated in HCM and RCM but downregulated in DCM. Our analysis also indicated significant downregulation of fatty acid precursor, arachidonic acid metabolism. Surprisingly, it was downregulated in the HCM and RCM phenotypes but upregulated in the ACM and DCM phenotypes (see Figure 3). From our results, we concluded that DCM displayed opposite trends compared to the other primary cardiomyopathies in critical metabolic pathways, such as glycolysis/gluconeogenesis, oxidative phosphorylation, riboflavin, thiamine, and purine metabolism.

**Figure 3:**
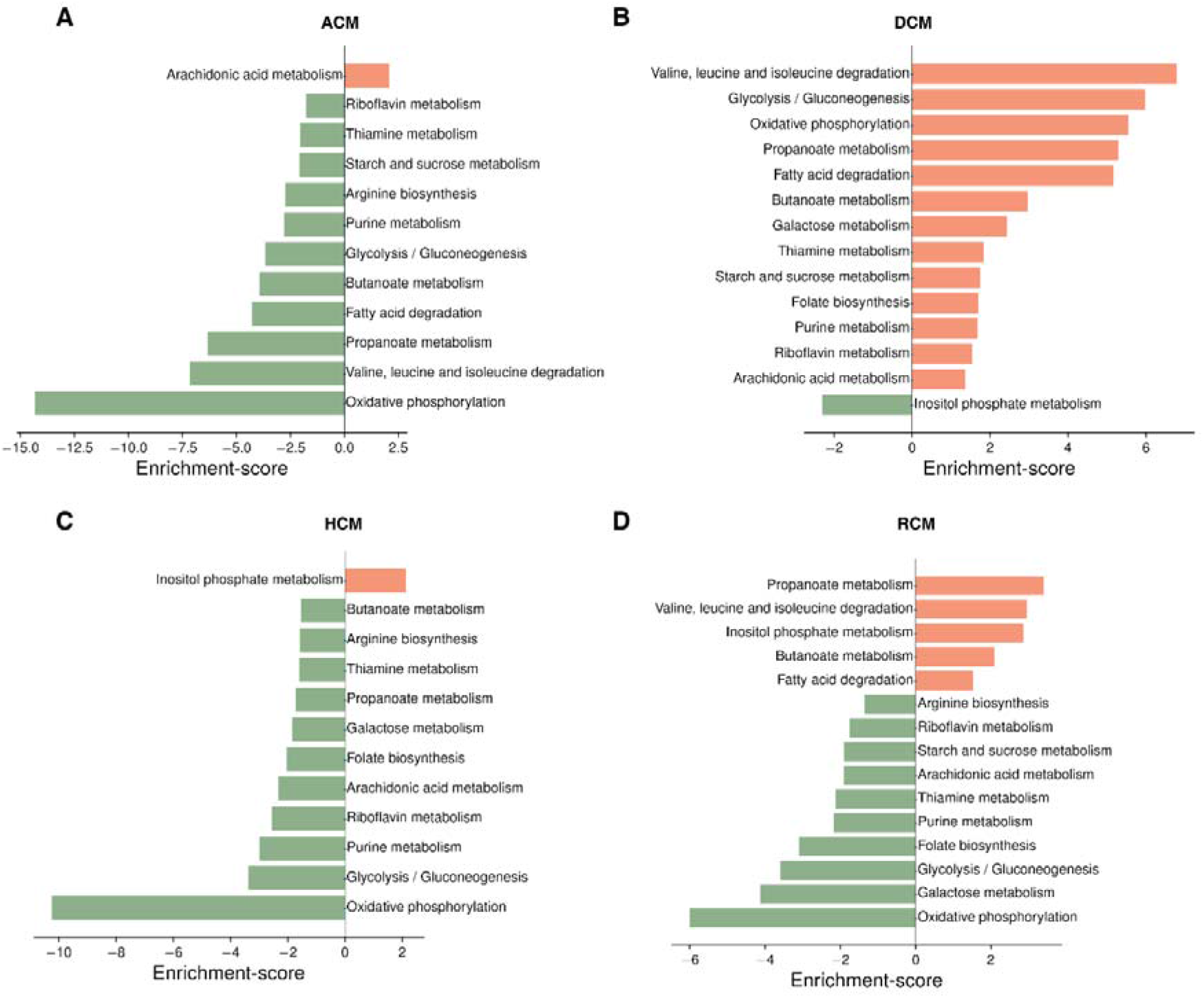
Top pathways enriched in ACM, DCM, HCM and RCM phenotypes (A,B,C,D).

### Arachidonic Acid Metabolism is Perturbed in Primary Cardiomyopathies

From the previous results, we focused on arachidonic acid metabolism due to its involvement in all cardiomyopathies and its less studied nature in cardiomyopathies. We chose independent studies from microarray datasets to validate AA metabolism perturbation in the ACM, DCM, and HCM human heart tissues. After GSA analysis on validation datasets, we observed that ~ 76% of KEGG pathways enriched in the original RNA-seq datasets were also significantly enriched in microarray sets. In the validation set, too, AA was dysregulated across ACM, DCM and HCM phenotypes (see Supplementary Figure S1). We focused on DEGs to further understand molecular players in AA metabolism. We identified 25 AA metabolism DEGs in primary cardiomyopathies (see Table 1). Genes like *AKR1C3, CYP2J2, EPHX2, LTC4S, PLA2G2A, PLA2G5, PTGDS* and *PTGIS* were particularly interesting (see Figure 4 and Supplementary Figure S2). The aldo/keto reductase superfamily protein-coding gene *AKR1C3* was downregulated in DCM and HCM but upregulated in ACM. The cytochrome P450 superfamily protein-coding gene *CYP2J2* was upregulated in ACM, DCM and HCM phenotypes. The epoxide hydrolase family protein-coding gene *EPHX2* was upregulated in DCM and RCM but downregulated in ACM. The MAPEG family protein-coding gene *LTC4S* was downregulated in HCM while upregulated in ACM and DCM. The phospholipase A2 family (PLA2) protein-coding gene *PLA2G2A* was upregulated in DCM and downregulated in HCM and RCM. At the same time, the PLA2 family protein-coding gene *PLA2G5* and glutathione-independent prostaglandin D synthase enzyme coding-protein *PTGDS* were downregulated in ACM, HCM and RCM. Lastly, the cytochrome P450 superfamily protein-coding gene *PTGIS* was upregulated in ACM, while downregulated in DCM and HCM (see Figure 5A). In summary, AA genes are significantly dysregulated in primary cardiomyopathies.

**Table 1:** A list of differentially expressed genes in arachidonic acid (AA) metabolism in arrhythmogenic, dilated, hypertrophic and restrictive cardiomyopathies.

| Gene | AA metabolism DEGs (log2FC) |  |  |  |
| --- | --- | --- | --- | --- |
|  | ACM | DCM | HCM | RCM |
| AKR1C3 | 0.7539 | -0.9029 | -0.324 | NA |
| ALOX5 | 0.7306 | NA | NA | NA |
| CBR1 | NA | 0.5106 | NA | NA |
| CBR3 | 0.6172 | NA | NA | NA |
| CYP2J2 | NA | 1.669 | 0.5889 | NA |
| CYP2U1 | 0.3272 | -0.6522 | 0.3382 | NA |
| EPHX2 | -0.4213 | 0.7662 | NA | 0.628 |
| GGT1 | NA | 1.1862 | NA | -0.8778 |
| GGT5 | 1.0935 | NA | -0.4816 | NA |
| GPX1 | NA | NA | -0.4593 | NA |
| GPX3 | NA | 1.2371 | -0.4969 | NA |
| GPX7 | 0.3851 | NA | NA | NA |
| GPX8 | 0.6545 | NA | NA | NA |
| LTC4S | 0.566 | 1.174 | -0.6461 | NA |
| PLA2G2A | NA | 2.1743 | -1.3455 | -1.1266 |
| PLA2G4A | 0.4587 | NA | NA | NA |
| PLA2G4C | NA | 0.8315 | NA | NA |
| PLA2G5 | -0.2836 | NA | -0.3504 | -0.5976 |
| PLA2G6 | 0.4439 | -0.6529 | NA | NA |
| PTGDS | -1.0987 | NA | -0.5926 | -0.9405 |
| PTGES | NA | NA | NA | -1.3769 |
| PTGES2 | NA | 1.2104 | NA | NA |
| PTGES3 | -0.3304 | 0.4085 | NA | NA |
| PTGIS | 0.7085 | -1.3175 | -0.8043 | NA |
| TBXAS1 | 0.4671 | NA | -0.9969 | NA |

**Figure 4:**
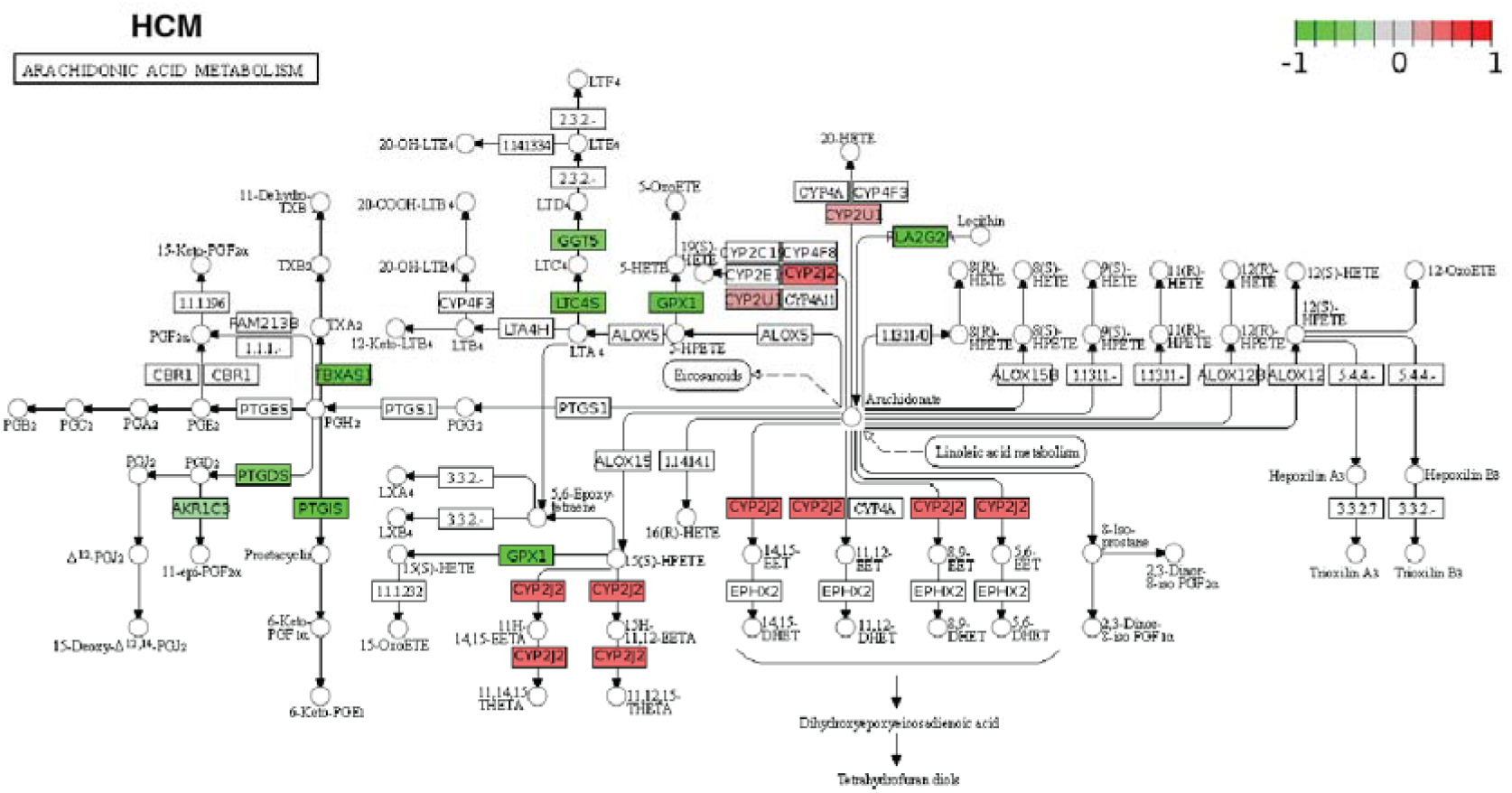
The KEGG pathway map of AA metabolism DEGs.

**Figure 5:**
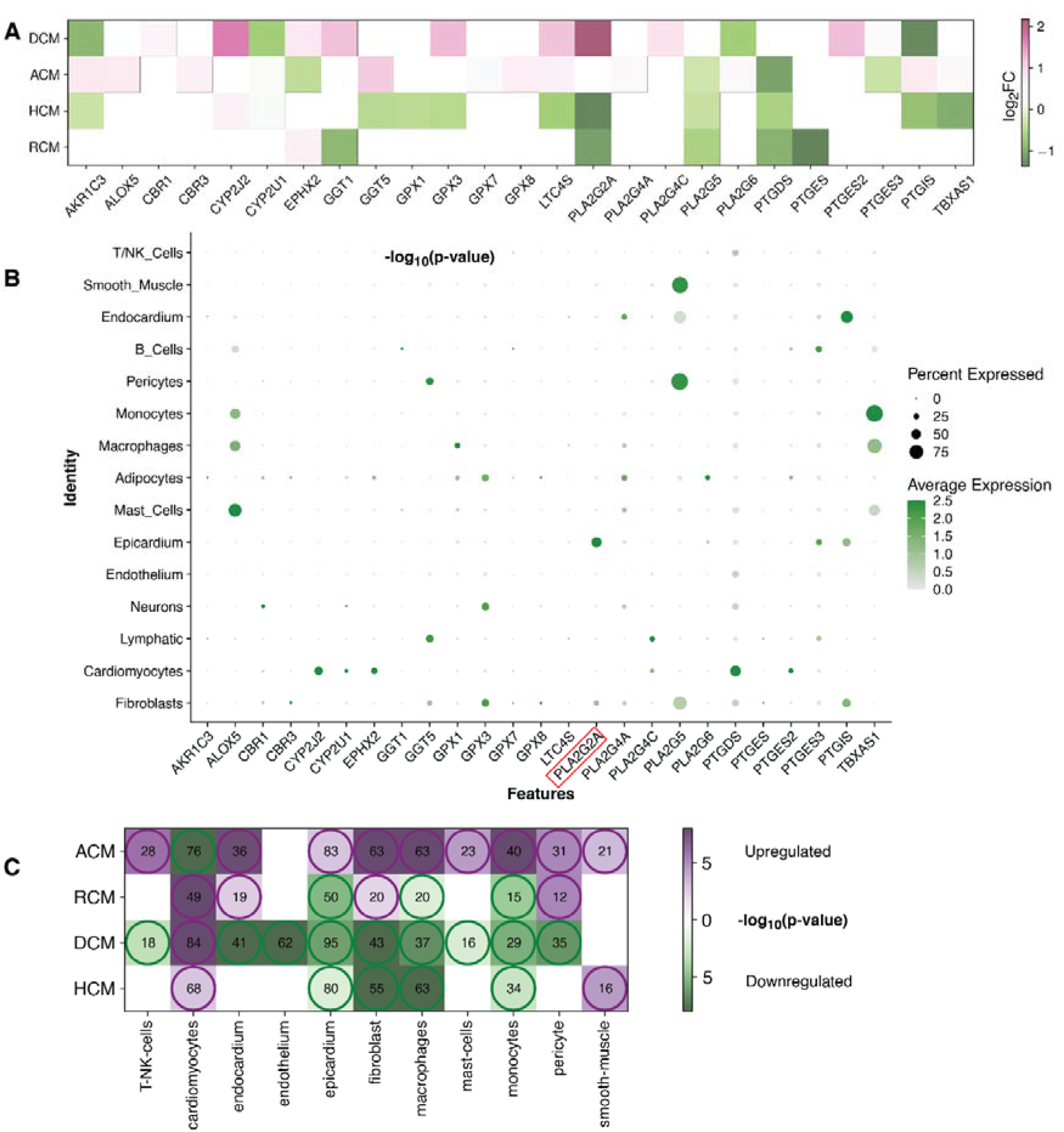
AA DEGs genes and their expression heart cell types. (A) A heatmap showing all AA DEGs in each cardiomyopathy (ACM, DCM, HCM and RCM). (B) The bubble plot showing expression of AA metabolism DEGs in heart tissue sn-RNA Seg dataset. (C) A Heatmap showing the cell type marker genes enriched DEGs in ACM, DCM, HCM and RCM phenotypes.

### Arachidonic Acid Metabolism Regulators Cell Types and Marker Genes in These Cells

Next, we performed meta analysis of the expression of the previous 25 AA metabolism genes in heart cell types. We utilized the cell types annotated in the human heart from an earlier single-nucleus RNA sequencing (snRNA-seq) study (GSE183852) (33). These cells were grouped into 15 clusters. Earlier mentioned genes, *AKR1C3, CYP2J2, EPHX2, LTC4S, PLA2G2A, PLA2G5, PTGDS* and *PTGIS*, were predominantly expressed in cardiomyocyte, epicardium, smooth muscle, endocardium, pericytes and fibroblasts (see Figure 5B). Interestingly, genes common in up to two phenotypes were expressed in more immune cells like mast cells, macrophages and monocytes. The lipoxygenase gene family member *ALOX5*, the glutathione peroxidase family member *GPX1* and the cytochrome P450 superfamily member *TBXAS1* belong to this category. The glutathione peroxidase family member *GPX3, PLA2G5* and *PTGIS* were expressed in a significant number of fibroblasts (greater than 50%). The presence of *AKR1C3* and *LTC4S* was negligible in all the major cell types of the heart.

T/NK cells, cardiomyocytes, endocardium, endothelium, epicardium, fibroblast, macrophages, mast cells, myocytes, pericytes and smooth muscles were further explored for marker genes identification. Few AA metabolic genes were present in multiple cell types but were left out as marker genes in all cell types as per the marker definition strategy. *PLA2G2A* gene is such an example that was expressed in fibroblast and epicardium. However, due to its higher expression in the epicardium, it was missed as a marker in fibroblast. Once the marker genes in the above cell types were identified, GSA was carried out on these genes set to inquire enrichment of these markers within upregulated or downregulated genes of ACM, DCM, HCM and RCM (see Figure 5C, supplementary Table S2-S5). The cardiomyocytes marker genes were enriched within upregulated genes of DCM, HCM and RCM (84, 68, 49) but within downregulated genes of ACM (76) (enrichment *p-values* < 0.1 and gene *adjusted P* < 0.1 & *|log_2_FC|* ≥ 0.28). In comparison, marker genes of fibroblasts were enriched within upregulated genes of ACM and RCM (63, 20) and within downregulated genes of DCM and HCM phenotypes (enrichment *p-values* < 0.1 and gene *adjusted P* < 0.1 & *|log_2_FC|* ≥ 0.28). Moreover, the marker genes of pericyte and endocardium were enriched within upregulated genes of ACM and RCM but within downregulated genes of DCM. Likewise, the smooth muscle marker genes were enriched within upregulated genes of ACM and RCM only. In this, a total of 21 and 16 marker genes were found to be upregulated (enrichment *p-values* < 0.1 and gene *adjusted P* < 0.1 & *|log_2_FC|* ≥ 0.28). The marker genes of the epicardium, macrophages and monocyte were enriched within downregulated genes of DCM, HCM and RCM and opposite in ACM phenotype. The T/NK cells and mast cell marker genes were enriched in the upregulated genes of ACM but within downregulated genes of DCM. Lastly, endothelium marker genes were enriched within upregulated genes of DCM (62) only (enrichment *p-values* < 0.1 and gene *adjusted P* < 0.1 & *|log_2_FC|* ≥ 0.28). Overall, expression of AA genes in heart tissue cell types and enrichment analysis on marker genes of these cell types demonstrates that AA metabolic genes are expressed in cardiomyocytes, fibroblast and immune cells, and a large proportion of cell type marker genes are dysregulated in all primary cardi omyopathi es.

### Network Analysis Reveals Association of Key Arachidonic Acid Metabolism Regulator *PLA2G2A* and Fibroblast Marker Genes

We next performed in-depth analysis of the significantly enriched marker genes within DEGs of primary cardiomyopathies. Gene ontology analysis of cardiomyocytes and fibroblast cell type enriched marker genes revealed biological processes associated with muscle contraction and extracellular matrix organization, confirming the relevance of these markers (see Supplementary Figure S3-S5). The AA metabolism DEGs and cell type enriched marker genes were mapped to human interactome to understand their underlying association (see Methods). Cardiac fibrosis is a significant player in cardiomyopathies. Therefore, we examined fibroblast cell types in this analysis. The fibroblast cell type dysregulated marker genes were associated with the *PLA2G2A* gene of AA metabolism in ACM, DCM and HCM (see Figure 6 and Supplementary Figure S6-S8). In the RCM phenotype, the interaction was missing, possibly due to much fewer overall DEGs in RCM. Network analysis of dysregulated HCM marker genes revealed that AA gene *PLA2G2A* interacts with *DCN*, which interacts with *FN1, COL4A4, SLIT2, EGFR GSN, COL1A2, ROBO1, COL4A1* and *ELN* that are primarily involved in extracellular matrix (ECM) organization and heart development. Similarly, DCM specific dysregulated marker genes network uncovered *PLA2G2A* interaction with *DCN*, which was directly linked to *FBN1* and *COL4A1*. These genes are primarily involved in ECM organization and anatomical structure morphogenesis. Collectively, network analysis demonstrates that the phospholipase A2 family gene *PLA2G2A* influences fibroblast and may be involved in fibrosis during cardiomyopathy.

**Figure 6:**
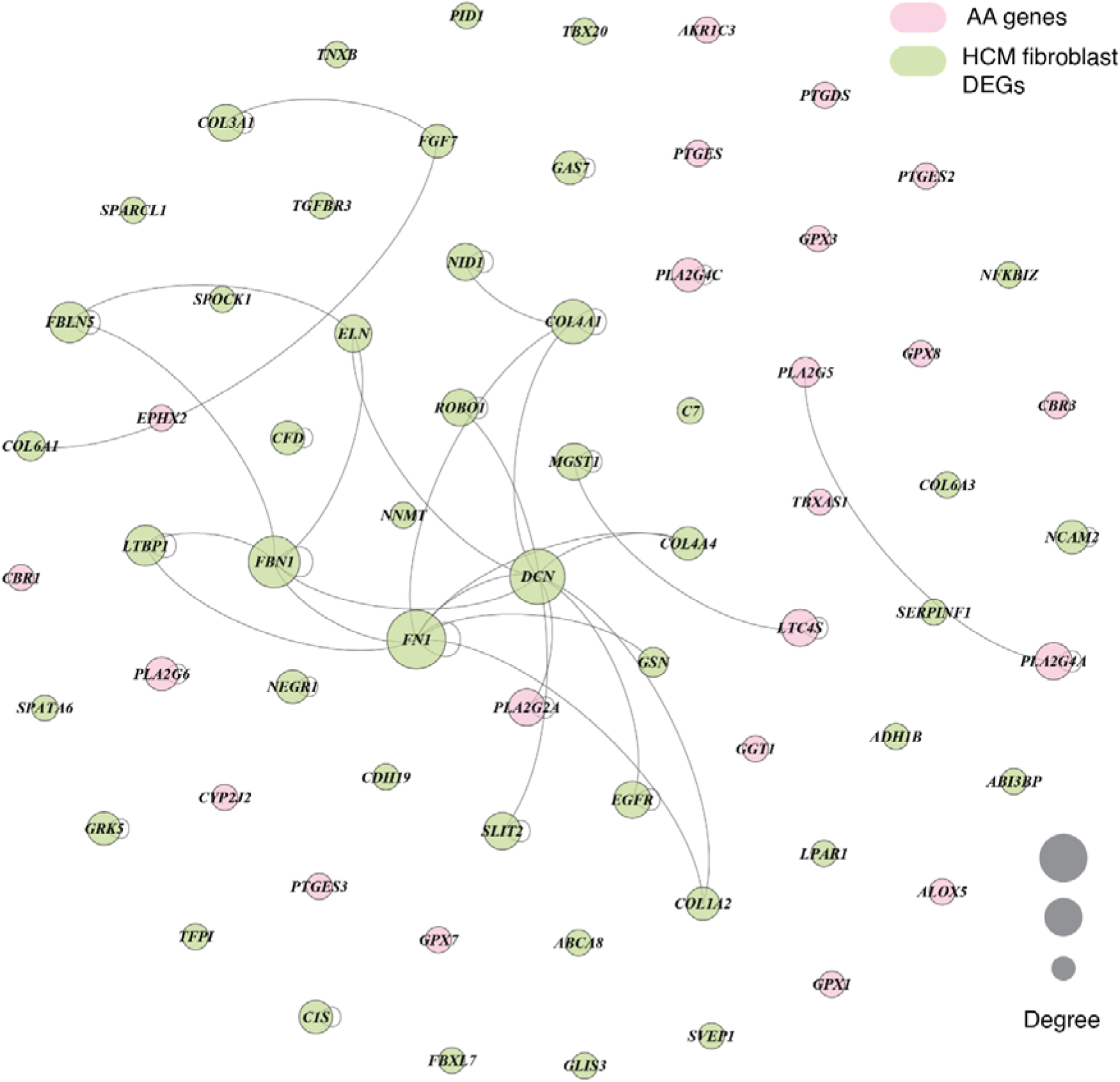
Network showing the interaction between AA metabolism DEGs and cell type marker DEGs in fibroblast cell type in HCM phenotype.

## Discussion

Primary cardiomyopathies like ACM, DCM, HCM and RCM are heterogenous heart muscle diseases with poor prognosis and a leading cause of sudden cardiac death (6–12). The current study accomplishes GSA on the transcriptome profile to understand metabolic pathways perturbation across primary cardiomyopathies. Among the GSA-enriched KEGG metabolic pathways between cardiomyopathy and normal samples, glycolysis/gluconeogenesis, oxidative phosphorylation, nucleotides metabolism, amino acid biosynthesis, co-factor metabolism, and fatty acid precursor arachidonic acid metabolism were statistically significant. Previous studies have reported oxidative phosphorylation, glycolysis and fatty acid metabolism perturbation in individual cardiomyopathies (23,24). In this work, we carried out an integrated analysis of AA metabolic perturbation across cardiomyopathies.

Arachidonic acid (AA) metabolism is an important mediator of cardiovascular processes such as fibrosis and inflammation. However, its role in cardiomyopathies is less explored (31,32). Our study highlighted the statistically significant dysregulation of AA metabolism in all primary cardiomyopathies. It was also consistent in the validation set. Further, AA metabolic DEGs in our study were expressed in heart tissue cell types like cardiomyocytes, fibroblasts, epicardium, endocardium, smooth muscle, and pericytes and immune cells such as macrophages, monocytes and mast cells at the single-cell level. These results point to a potential role of AA metabolism in modulating fibrosis and inflammation in cardi omyopathi es.

Cardiomyopathies are generally characterized by cardiac fibrosis (44). Cardiac fibrosis results from dysregulation of the balance between the synthesis and degradation of extracellular matrix (ECM) proteins (44,45). Fibroblast cells synthesize the ECM proteins and get involved in tissue repair and remodelling after injury in heart tissues during cardiomyopathies (44–46). We investigated if the AA metabolism genes were associated with dysregulated fibroblast marker genes. Our analysis showed the interaction between the AA metabolism gene *PLA2G2A* and fibroblast marker gene *DCN* that in turn, was linked to many other fibroblast marker genes. The gene ontology enrichment analysis of fibroblast marker genes confirmed their role in ECM organization. The decorin (*DCN*) belongs to chondroitin sulfate proteoglycan and has been shown to exhibit antifibrotic effects in the mouse model (47). In short, our analysis demonstrated that AA metabolism genes interact with the ECM proteins, such as decorin and others, directly or indirectly and might influence cardiac fibrosis in primary cardiomyopathies.

This study has limitations due to heterogeneity in samples aggregated from different transcriptome studies. Sequencing methodology, environment and sequencing depth are the main challenges in an integrative analysis. Quantification of RCM RNA-seq reads was missing for many genes due to less coverage of these genes. Next, this study only demonstrated statistically perturbed pathways, important DEGs and heart cell types and their interactions in each cardiomyopathy through computational analysis. The experimental validation in an in vitro or in vivo system was not performed due to the limitation of resources. Lastly, ACM, DCM, HCM and RCM are highly heterogeneous diseases. Thus, the classification of these may vary substantially in the population.

Regardless of the limitations, the current study provides new insights into cardiomyopathy research. This work recognises the arachidonic acid metabolism as a potential regulator of all major primary cardiomyopathies. Apart from this, the study also demonstrates the expression of AA metabolism genes in major heart-specific cell types and immune cells, facilitating a better interpretation of its potential roles in the disease. Interaction between AA gene *PLA2G2A* and fibroblast marker gene *DCN* have a potential role as a therapeutic target to regulate cardiac fibrosis.

## Conclusion

Arachidonic acid metabolism is perturbed in primary cardiomyopathies. AA metabolism genes are expressed in most heart cell types and immune cells, influencing the immune activation and cardiac fibrosis. The association of dysregulated AA gene *PLA2G2A* with fibroblast marker gene *DCN* may be an important factor related to fibrosis.

## Data availability statement

This paper presents an analysis of existing, publicly available data. All codes used in this study can be found on the following link http://caps.ncbs.res.in/download/cardiomyo_transcriptome.

## Author Contributions

RS and PKC conceptualized the study. PKC integrated data, performed data analysis and visualization. PKC wrote the manuscript and RS wrote the final draft of manuscript.

## Acknowledgments

We acknowledge the contributions from the publicly available databases as source of original data in our study. We want to thank CAPS lab members for their technical help and discussions. This work was inspired by the heart ailments of our collaborator, late Prof N. Srinivasan of Indian Institute of Science. The authors thank NCBS (TIFR) for infrastructural facilities. PKC would like to thank NCBS-TIFR (Department of Atomic Energy, India) for the fellowship.

## Supplementary material

The supplemental information for this study is available in the Supplementary file.

