## Supplemental Information for "Transcriptome Data Analysis of Primary Cardiomyopathies Reveals Perturbations in Arachidonic Acid Metabolism"

**Supplementary Figures and Tables**

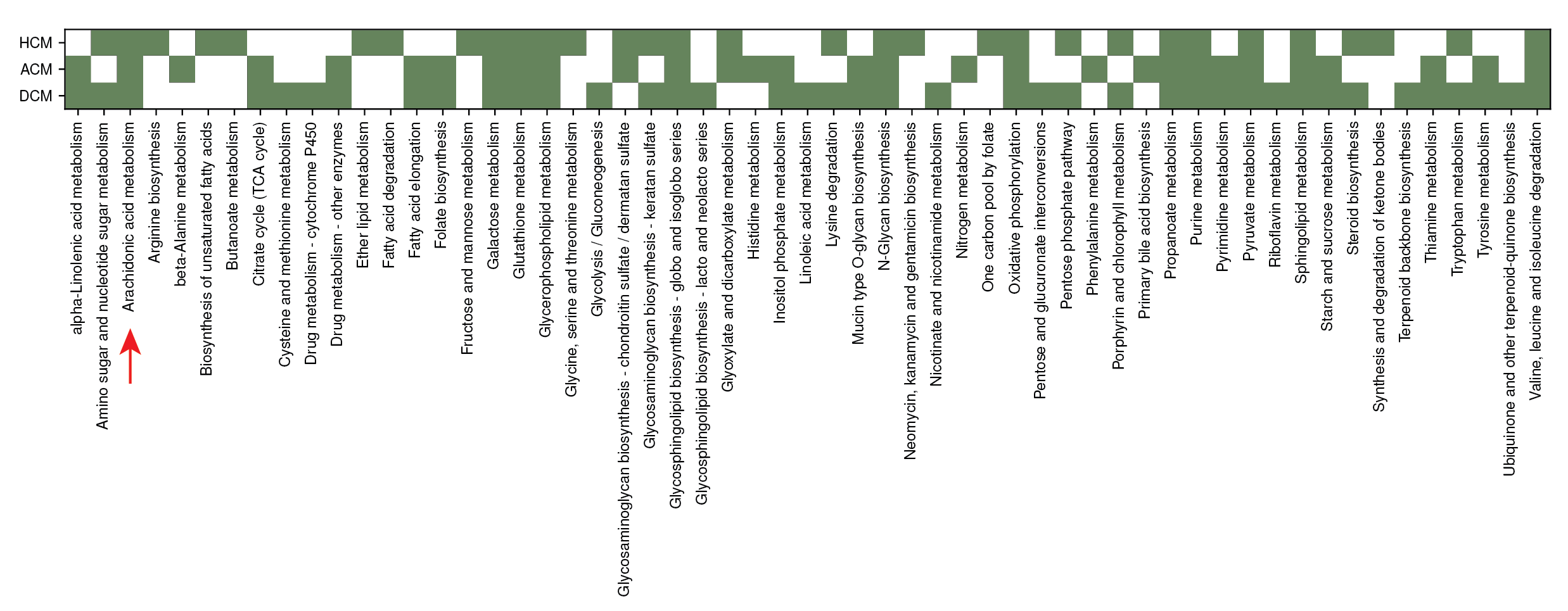

**Figure S1: Perturbation of major pathways in hypertrophic cardiomyopathy, arrhythmogenic cardiomyopathy and dilated cardiomyopathy**

This figure shows heatmap of statistically significant pathways from GSA analysis on KEGG pathways in HCM, ACM and HCM microarray studies.

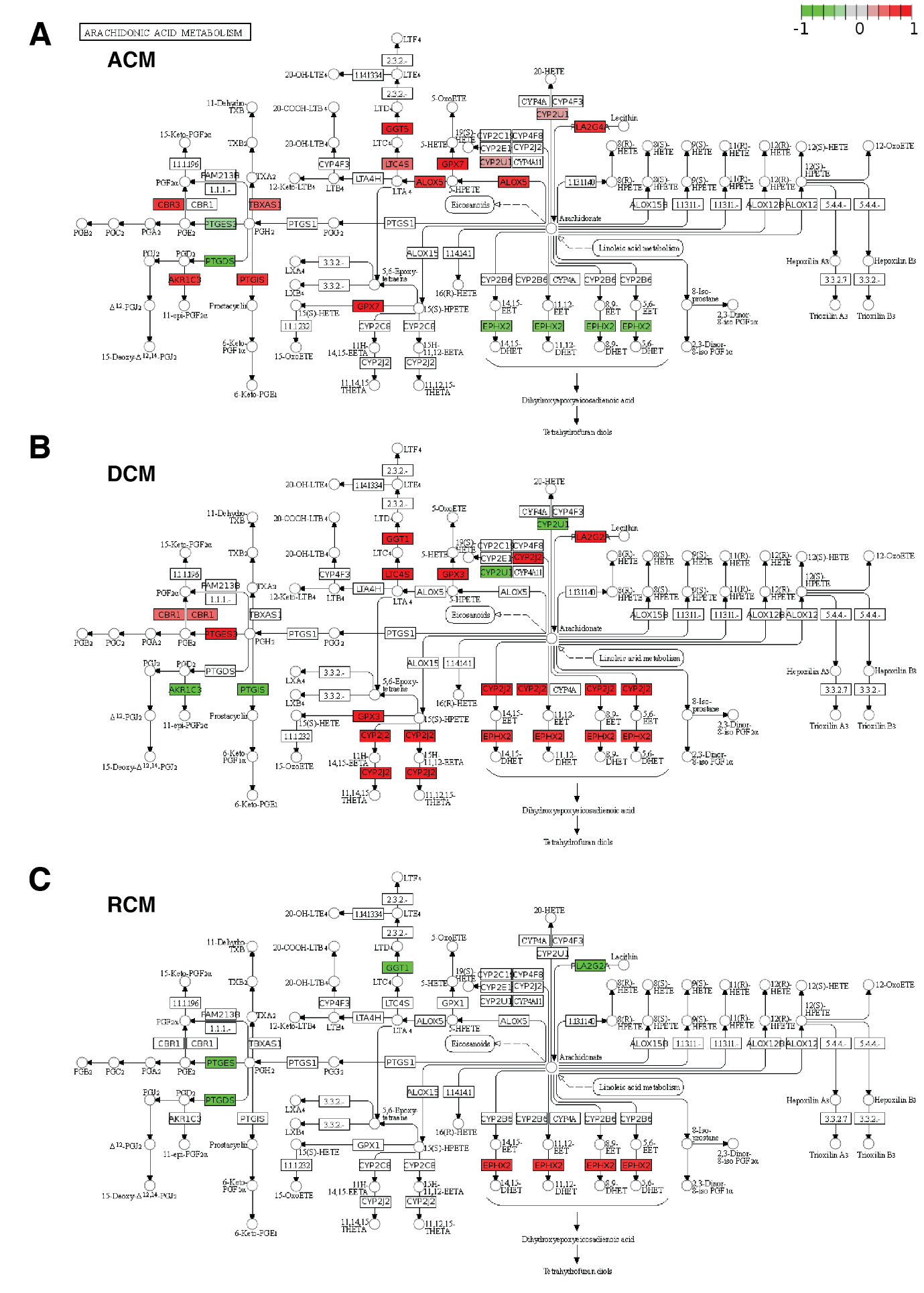

**Figure S2: Arachidonic acid (AA) metabolism DEGs**

Graph illustration of arachidonic acid metabolism DEGs on the KEGG pathway map. (**A,B,C**) Panels correspond to significantly dysregulated AA metabolism genes (*adjusted P* < 0.1 and |*log_2_FC|* ≥ 0.28) in arrhythmogenic cardiomyopathy (**A**), dilated cardiomyopathy (**B**), and restrictive cardiomyopathy (**C**), respectively.

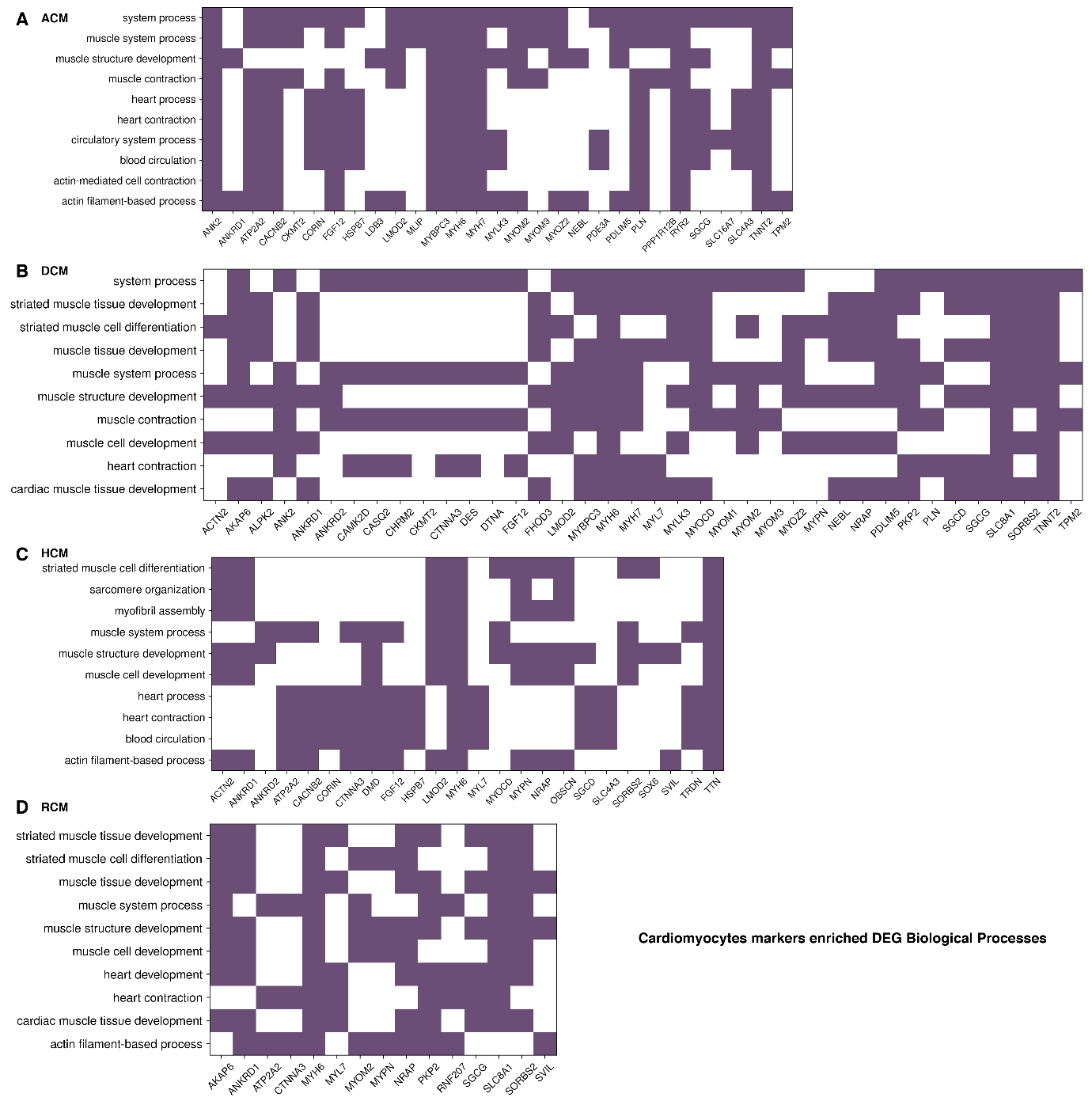

**Figure S3: Functional enrichment of cardiomyocytes specific marker DEGs**

Heatmap visualization of biological processes associated with cardiomyocytes markers DEGs in ACM (**A**), DCM (**B**), HCM (**C**), and RCM (**D**) datasets.

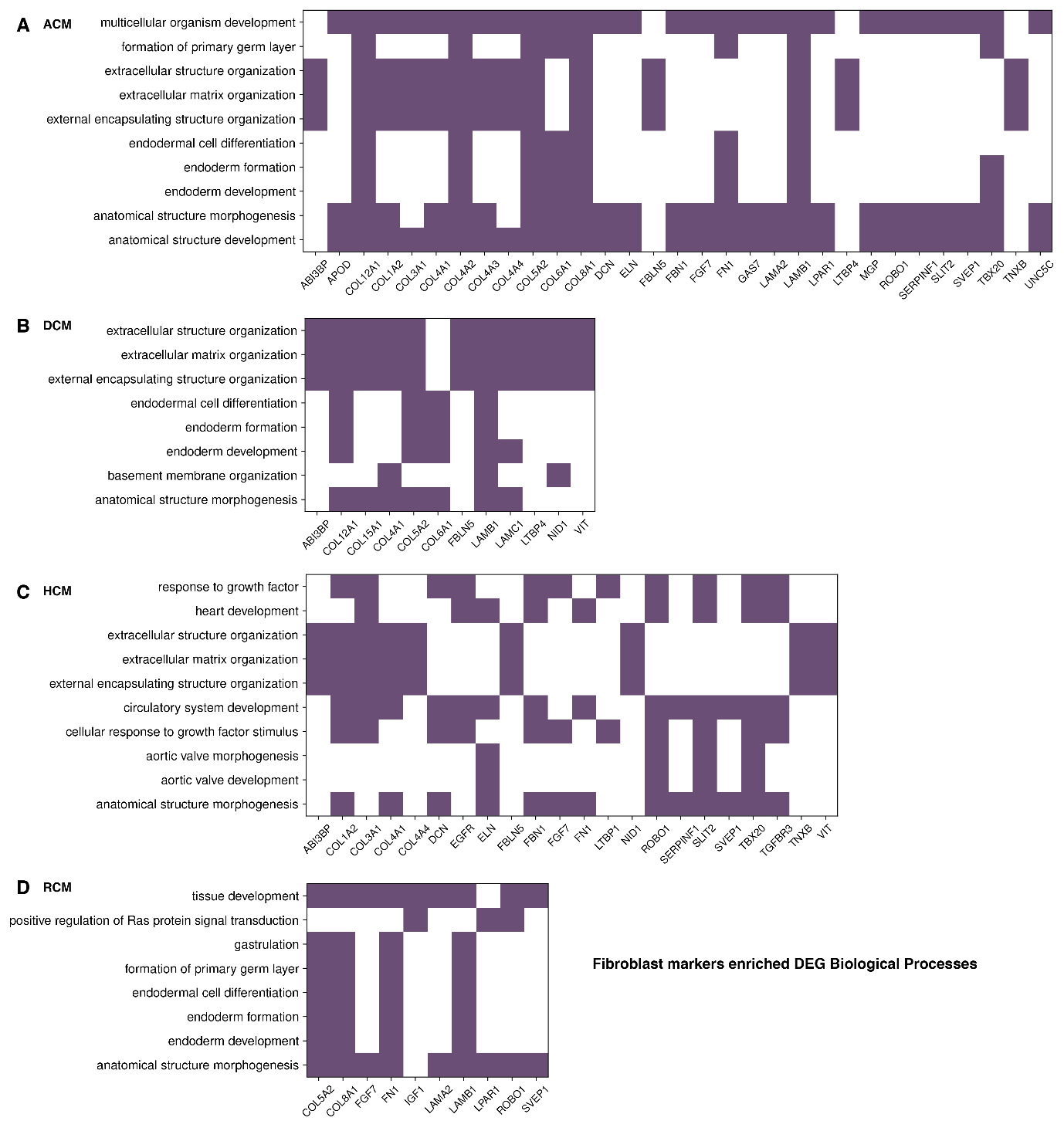

**Figure S4: Functional enrichment of fibroblasts specific marker DEGs**

Heatmap visualization of biological processes associated with fibroblasts markers DEGs in ACM (**A**), DCM (**B**), HCM (**C**), and RCM (**D**) datasets.

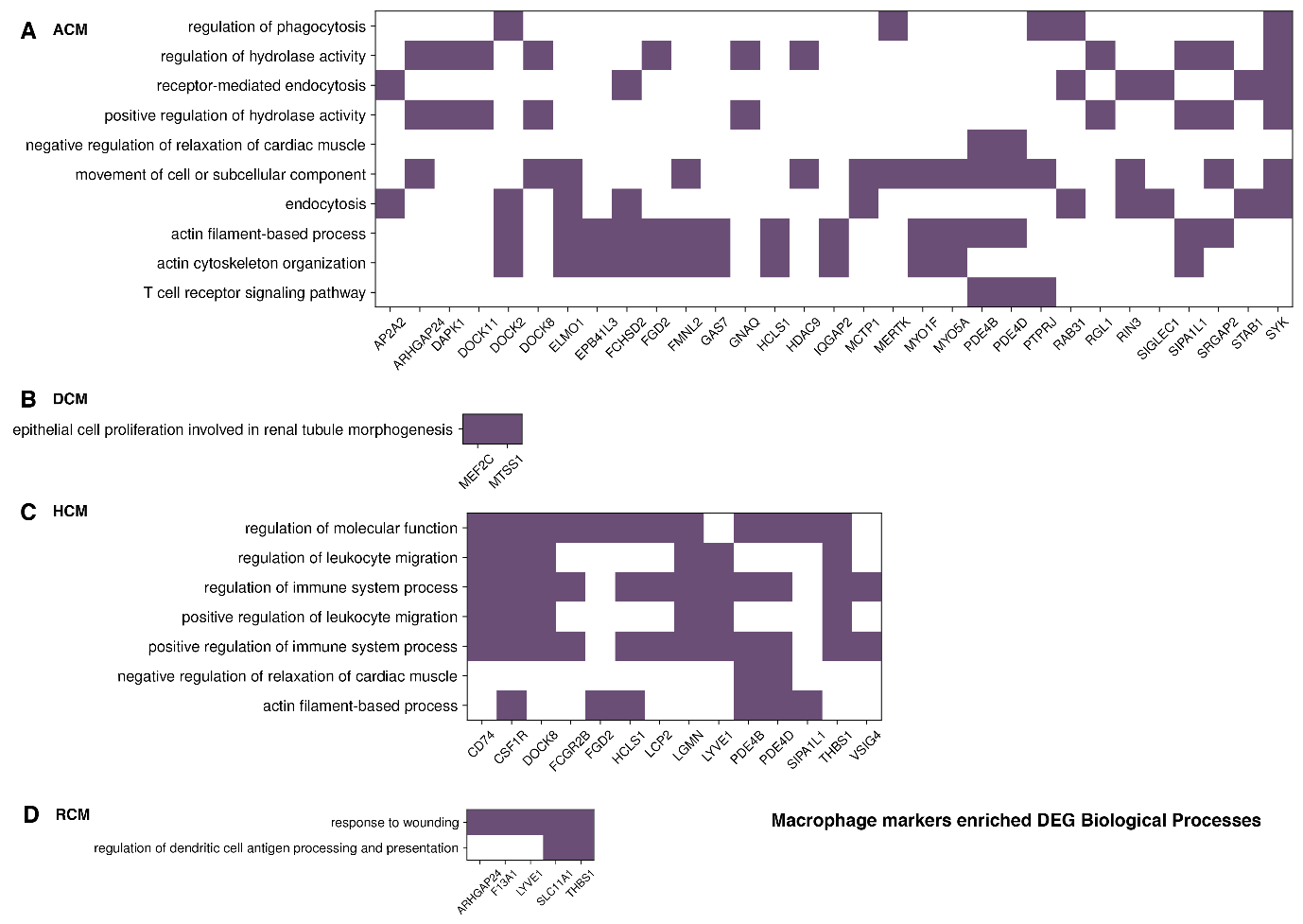

**Figure S5: Functional enrichment of macrophage specific marker DEGs**

Heatmap visualization of biological processes associated with macrophage markers DEGs in ACM (**A**), DCM (**B**), HCM (**C**), and RCM (**D**) datasets.

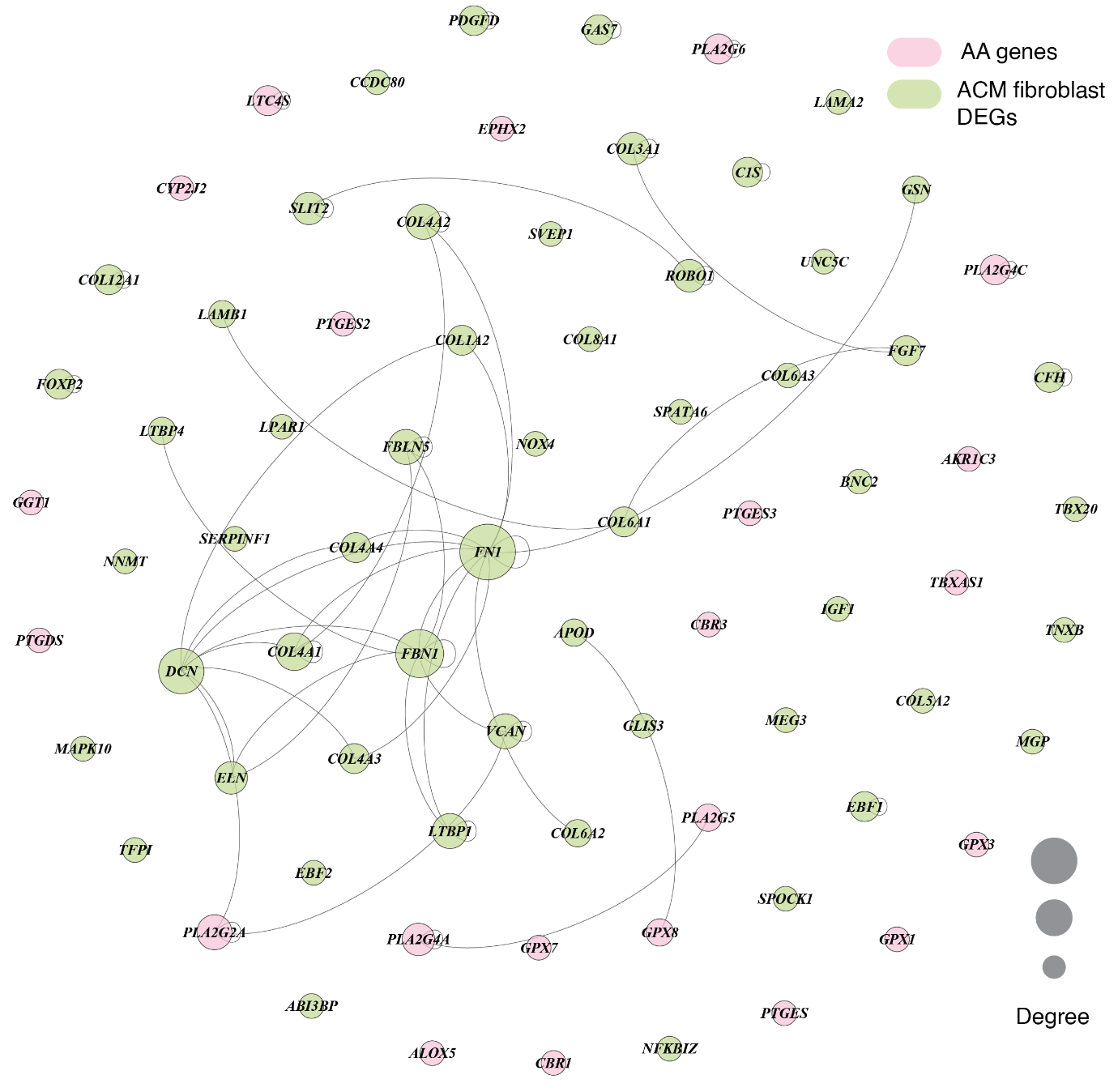

**Figure S6: Interaction network of dysregulated AA metabolism genes and ACM fibroblast marker DEGs on human interactome**

A network sub-graph showing the interaction between the differentially expressed AA metabolism genes and fibroblast marker DEGs in ACM phenotype on the human interactome. The node size in the network corresponds to the degree of each node. The AA metabolism genes are coloured in pink while marker genes are marked with green colour in the sub-graph.

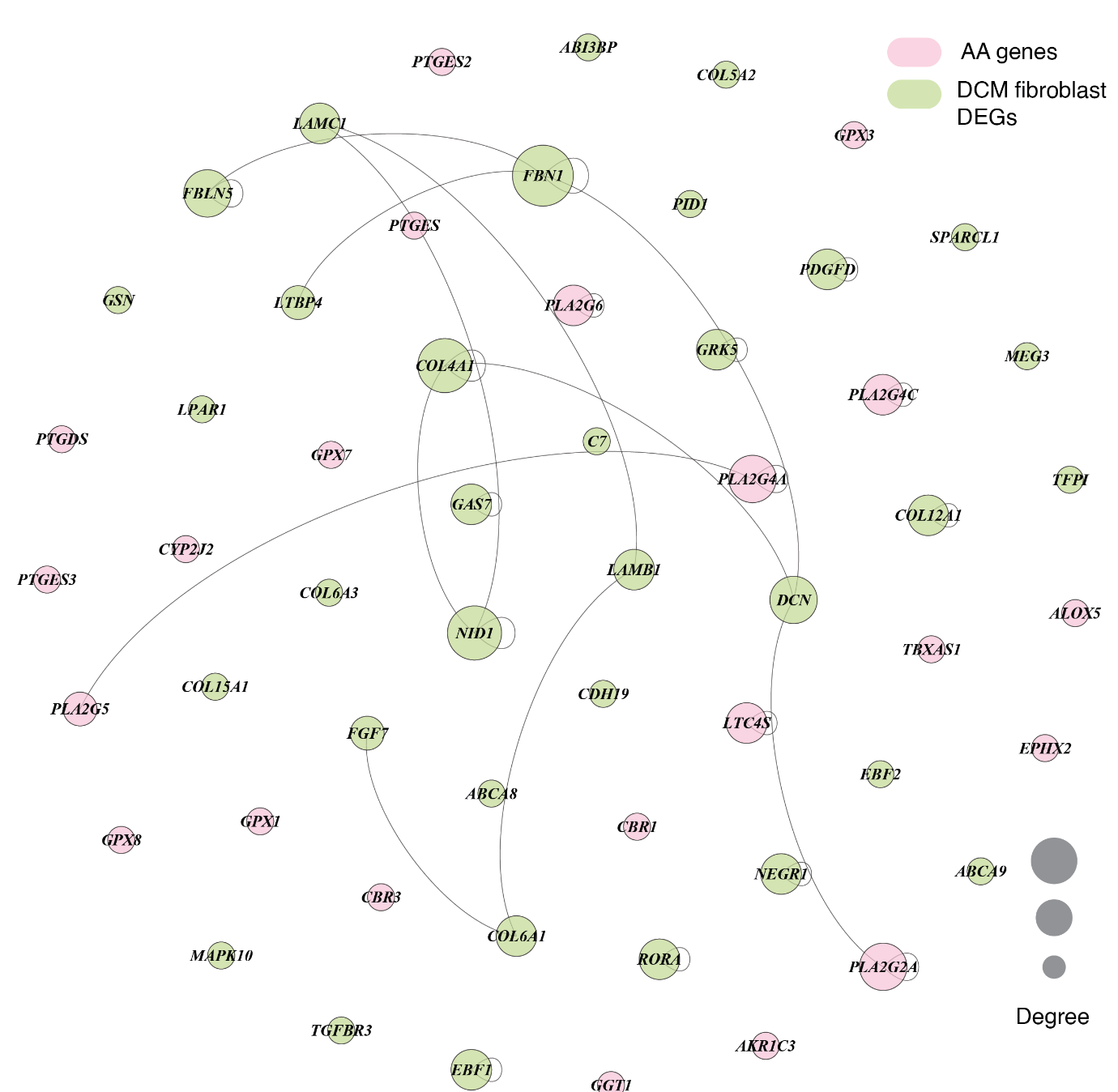

**Figure S7: Interaction network of dysregulated AA metabolism genes and DCM fibroblast marker DEGs on human interactome**

A network sub-graph showing the interaction between the differentially expressed AA metabolism genes and fibroblast marker DEGs in DCM phenotype on the human interactome. The node size in the network corresponds to the degree of each node. The AA metabolism genes are coloured in pink while marker genes are marked with green colour in the sub-graph.

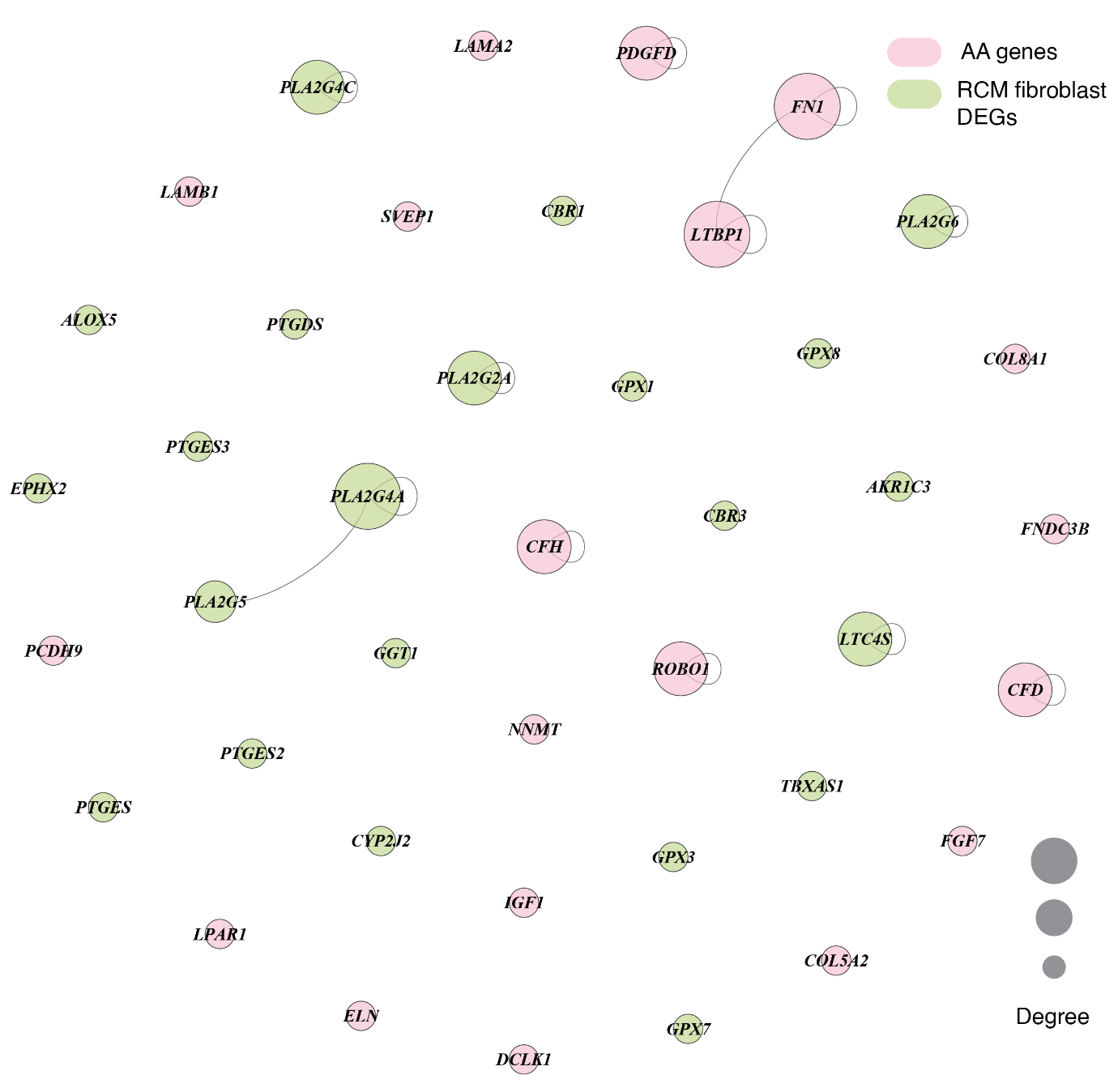

**Figure S8: Interaction network of dysregulated AA metabolism genes and RCM fibroblast marker DEGs on human interactome**

A network sub-graph showing the interaction between the differentially expressed AA metabolism genes and fibroblast marker DEGs in the RCM phenotype on human interactome. The node size in the network corresponds to the degree of each node. The AA metabolism genes are coloured in pink while marker genes are marked with green colour in the sub-graph.

**Table S1: Transcriptome data of ACM, DCM, HCM, and RCM phenotypes**

List of all transcriptome datasets considered in the study

**Table S2: Cell type marker DEGs in arrhythmogenic cardiomyopathy**

List of all dysregulated markers in enriched heart cell types in ACM. The cut-off value of *adjusted P* < 0.1 and |*log_2_FC|* ≥ 0.28 was used.

**Table S3: Cell type marker DEGs in dilated cardiomyopathy**

List of all dysregulated markers in enriched heart cell types in DCM. The cut-off value of *adjusted P* < 0.1 and |*log_2_FC|* ≥ 0.28 was used.

**Table S4: Cell type marker DEGs in hypertrophic cardiomyopathy**

List of all dysregulated markers in enriched heart cell types in HCM. The cut-off value of *adjusted P* < 0.1 and |*log_2_FC|* ≥ 0.28 was used.

**Table S5: Cell type marker DEGs in restrictive cardiomyopathy**

List of all dysregulated markers in enriched heart cell types in RCM. The cut-off value of *adjusted P* < 0.1 and |*log_2_FC|* ≥ 0.28 was used.
